# Powerful and interpretable behavioural features for quantitative phenotyping of *C. elegans*

**DOI:** 10.1101/389023

**Authors:** Avelino Javer, Lidia Ripoll-Sanchez, André E.X. Brown

**Affiliations:** MRC London Institute of Medical Sciences, London, UK; Institute of Clinical Sciences, Imperial College London, London, UK

**Keywords:** Computational ethology, C. elegans, Phenotyping, Worm tracking

## Abstract

Behaviour is a sensitive and integrative readout of nervous system function and therefore an attractive measure for assessing the effects of mutation or drug treatment on animals. Video data provides a rich but high-dimensional representation of behaviour and so the first step of analysis is often some form of tracking and feature extraction to reduce dimensionality while maintaining relevant information. Modern machine learning methods are powerful but notoriously difficult to interpret, while handcrafted features are interpretable but do not always perform as well. Here we report a new set of handcrafted features to compactly quantify *C. elegans* behaviour. The features are designed to be interpretable but to capture as much of the phenotypic differences between worms as possible. We show that the full feature set is more powerful than a previously defined feature set in classifying mutant strains. We then use a combination of automated and manual feature selection to define a core set of interpretable features that still provides sufficient power to detect behavioural differences between mutant strains and the wild type. Finally, we apply the new features to detect time-resolved behavioural differences in a series of optogenetic experiments targeting different neural subsets.

## Introduction

Measuring phenotypes is essential in most areas of biology, but there are no rules that determine which aspects of a phenotype to focus on. This has led to calls for more exhaustive characterisations of phenotype under the umbrella term phenomics [1,2]. Imaging is well suited to phenomics because images can capture complex morphological differences and videos provide a natural extension to measure dynamics. However, the raw pixel intensities in images do not map directly to most quantities of interest and so an element of choice in representation remains. The success of deep learning approaches demonstrates the usefulness of automatically learned features on image analysis problems [3]. However, the tasks solved in deep learning have well defined objectives such as minimising cross entropy loss. When the objective is scientific understanding or hypothesis generation, high performance depends not just on accuracy but on interpretability and the nonlinear combination of many features through neural networks does not typically lead to highly interpretable features. Our objective is to find a middle ground using a range of interpretable features optimised to quantify *C. elegans* morphology and behaviour.

In *C. elegans*, phenotyping morphology and behaviour, both of which are readily captured using imaging, has a long history dating back to Brenner’s original paper describing the isolation of the first visible mutants [4]. Subsequent work by Croll *et al*. pioneered the quantitative analysis of nematode behaviour in *C. elegans* and other species [5–9]. A desire to increase throughput, sensitivity, and the ability to quantify multiple phenotypes from the same recording, have led to the development of many worm trackers over the intervening decades. Dusenbury made a tracker in 1983 that could track the centroid position of 25 worms in real time at 1 Hz [10]. The tracker was used to study oxygen and carbon dioxide responses and responses to a variety of chemicals [11].

The next generation of trackers were used to quantify speed [12] or the behavioural components of chemotaxis [13], still at relatively low resolution. High resolution single worm trackers were developed first simply to record single animals for long periods for subsequent manual annotation of egg laying [14], but were quickly adapted for use in high-dimensional quantitative phenotyping [15–19]. Throughput was increased by using multiple single-worm trackers in parallel [20]. At the same time, multi-worm trackers that tracked many animals at lower resolution were developed to increase throughput using a single camera [21,22]. Improvements in camera technology eventually led to the development of a high resolution multi-worm tracker that operates in real time and records worm outline and skeleton at 30 Hz [23]. New trackers continue to be developed for specific applications or with new features [24–33]. We have also recently developed a high resolution multi-worm tracker to store not just worm outline and skeleton but also worm pixels to achieve compression without losing information about worms and their surroundings [34]. Keeping a portion of the image data enables reanalysis using improved computer vision algorithms or manual annotation. In summary, there is no shortage of methods for collecting worm behaviour data and several options for quantifying behavioural phenotypes. Given the large set of possible approaches and features, a principled way of selecting useful features would be helpful.

In this paper, we introduce a set of handcrafted features that can be measured from single or multi-worm tracking data, provided there is sufficient resolution to quantify worm posture. The features are inspired by phenomics to be as exhaustive as possible, but with an explicit bias towards interpretability to support exploratory analyses, generate hypotheses, and guide mechanistic studies. We use a large database of videos of mutant worms and wild isolates to select feature subsets that balance explanatory power and interpretability. We also analyse a new set of optogenetics experiments and show that the same feature set can be used to find differences in behavioural dynamics that reveal a range of behavioural responses to optogenetic stimulation of different neural circuits in worms.

## Materials and Methods

The data from the mutants and wild-isolates are from two previously published studies and are available online from the OpenWorm Movement Database community page on Zenodo https://zenodo.org/communities/open-worm-movement-database/. As described previously, the worms in these videos were young adults recorded for 15 minutes crawling on agar on a patch of *E. coli* OP50 which serves as a food source [20,35]. Worms were allowed to habituate to the tracking plates for 30 minutes before recording.

The optogenetics experiments were performed on young adults on OP50 that were allowed to habituate for 30 minutes prior to recording. Worms were recorded for 7 minutes without perturbation and then stimulated with blue LEDs (peak intensity at 467 nm) with five 5-second pulses and one 90 second pulse, each separated by 60 seconds. Tracking plates were prepared with 300 μL OP50 liquid culture mixed with all-trans-retinal (ATR) dissolved in ethanol to a final plate concentration of 83 μM ATR (0.25 μl ATR each 300 μl of OP50).

Control plates were prepared identically but using 100% ethanol without ATR. Plates were left for 48h to dry with the lids on and stored for up to 5 days at 4°C. See Table 2 for a list of strains used in the optogenetics experiments.

All worms were segmented, tracked, and skeletonised using Tierpsy Tracker. Binaries, source code, and documentation are available at http://ver228.github.io/tierpsy-tracker/.

## Results

### Feature definition

For the initial parameterisation of the worm we focussed on defining features that cover as much of the range of observable phenotypes as possible while remaining interpretable. We identified a range of feature classes that we then characterised by defining multiple features for each class to ensure phenotypic breadth. Following previous efforts at hand-crafting features for quantifying *C. elegans* [15–17,20,23,36], the classes cover morphology, path, posture, and relative and absolute velocities. Interpretability is subjective and depends on the assessor’s background. As a rule of thumb, we considered more derived features—that is, features requiring more computational steps or longer algorithms to calculate—to be less interpretable. However, for the initial parametrisation, we erred on the side of including more features because if a feature is subsequently found to be important for detecting strain differences it could be worth sacrificing some interpretability.

The starting features are shown schematically in Fig. 1 and are described in more detail in the Supplementary Information.

**Figure 1:**
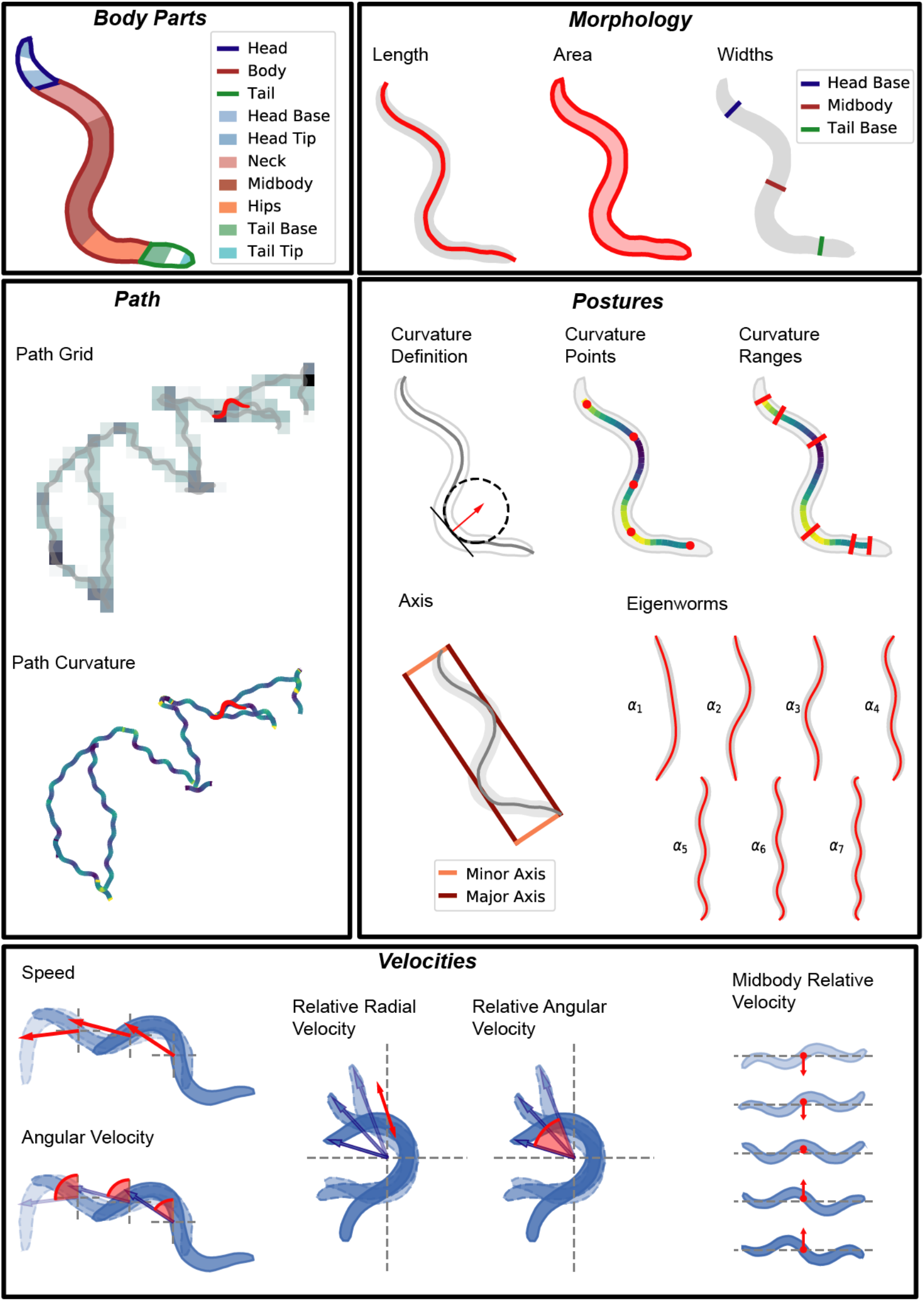
Schematics of the core features. Each of the features is summarized and expanded according to the transformations in Figure 2. See the supplementary information for detailed definitions.

### Feature expansion

To further increase the breadth of phenotyping, we perform a series of operations that expands the total number of features (Fig. 2). First, any feature that can be localised to a part of the body, for example curvature, is calculated separately for 5 segments along the worm (colloquially: head, neck, midbody, hips, and tail). Velocities are calculated additionally at the head tip and tail tip because motion at the extremities, especially the head, is often informative. Second, we calculate the derivatives of any time series features (i.e. features that are calculated in each frame). For example, we calculate the rate of change of curvature for each segment. Third, we subdivide features according to motion state (forward, backward, and pause) leading to features such as midbody curvature during reversals. Finally, the distributions of these features are quantified by calculating the 10^th^ percentile, median, and 90^th^ percentile values.

**Figure 2:**
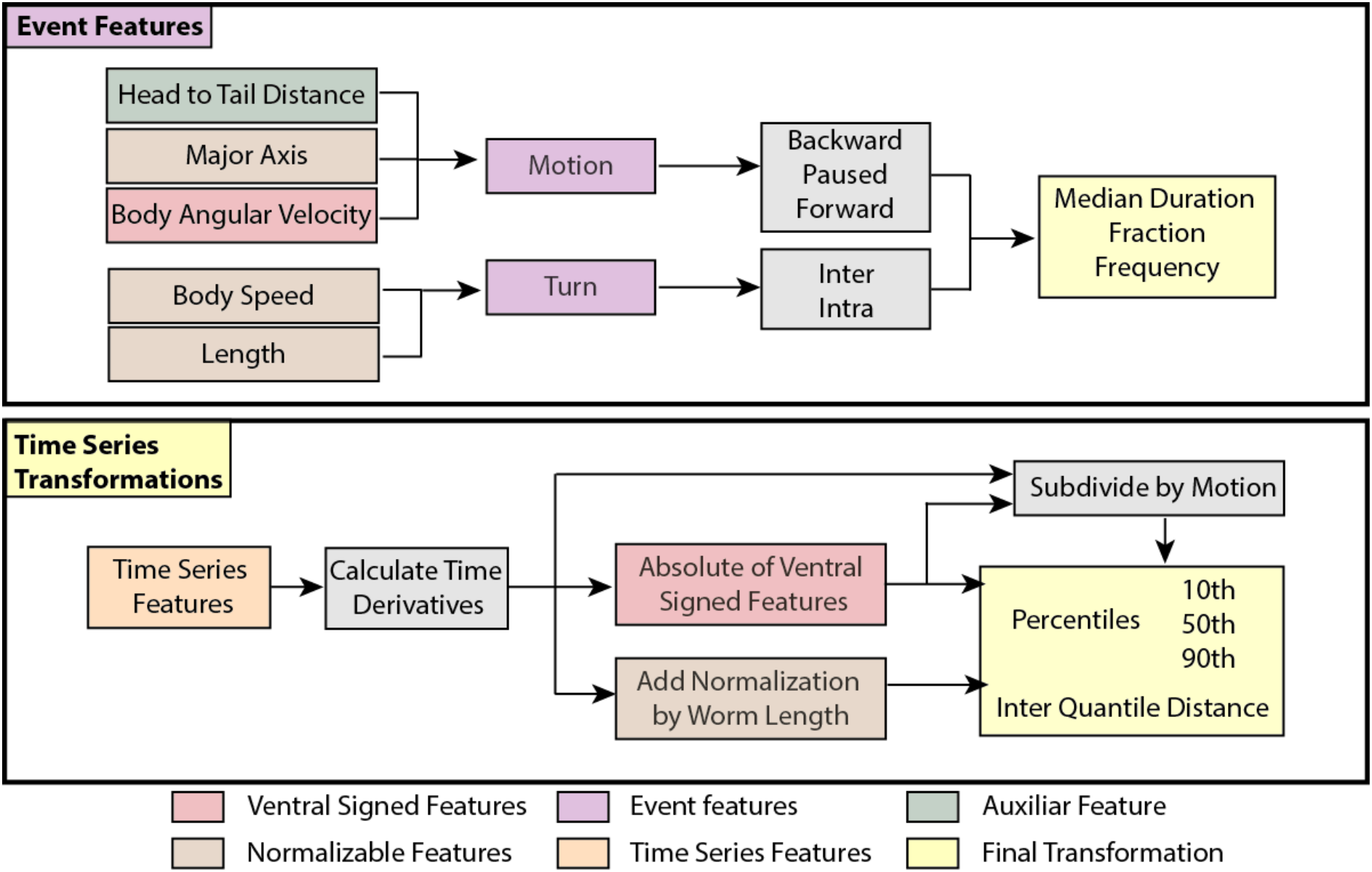
Operations that expand and summarize each of the core features. See Fig. S1 for a more detailed description on how the core time series features are subdivided.

The final phenotypic fingerprint thus derived has 4083 values. Although they are derived in several steps, they remain describable in words, for example, the feature *curvature_tail_w_forward_abs_90th* is the 90^th^ percentile of the absolute value of the tail curvature measured while the worm is moving forwards. The large number of features belies an underlying simplicity: there are only 16 basic features, each subjected to similar operations during expansion.

### Feature selection

Feature selection is useful to 1) remove noisy or irrelevant features and 2) to choose one from a set of highly correlated and therefore redundant features. Removing irrelevant features can improve performance while removing highly correlated features reduces the complexity of the representation without hurting performance. The features defined above involve relatively simple computations and were chosen based on our prior notions of what would be relevant for worm phenotyping so we do not expect many features to fall into the first category. On the other hand, the expansion procedure is likely to produce sets of correlated features that capture redundant information about the phenotype. In any case, relevance and redundancy are defined with respect to a particular dataset. We therefore chose to quantify the usefulness of the full set of features on a classification task on diverse previously published data sets [20,35] consisting of a total of 11,406 individual worms from 358 strains drawn from mutants affecting neurodevelopment, synaptic and extrasynaptic signalling, muscle function, and morphology as well as wild isolates representing some of the natural diversity of *C. elegans* strains around the world.

As a prepossessing step we cleaned the data by removing any feature assigned as NaN (Not a Number) in more than 2.5% of the worms in the full set. On this data set, only paused motion state features were eliminated because in 14.3% of the videos the worms never paused and therefore the subdivision is not defined. Any remaining NaN values are imputed using the population mean value of the corresponding feature. We then z-normalise the data by subtracting the feature mean and dividing by its standard deviation to make features with different units comparable on the same scale.

We divided the data into training and validation sets by randomly splitting the data per strain into 80%-20% for training and testing respectively. We then used recursive feature elimination to identify useful feature sets using the following procedure: 1) We fit a logistic regression model using stochastic gradient descent on the categorical cross entropy loss. 2) Each feature is ranked in importance by removing it from the fitted model and calculating the change in the loss. More important features will increase the loss when removed, while less important features will have little effect or even decrease the loss. 3) The least important features are dropped until the next power of two is reached, *e.g*. if there are 3000 features, 952 features will be dropped leaving 2048, or 2^11^. We repeated this procedure 10 times for different random subsets of worms and plotted the classification accuracy as a function of feature number for our newly defined features (Tierpsy features), the previously defined features from Yemini *et al*. [20], and the combination of both feature sets (Fig. 3a).

**Figure 3:**
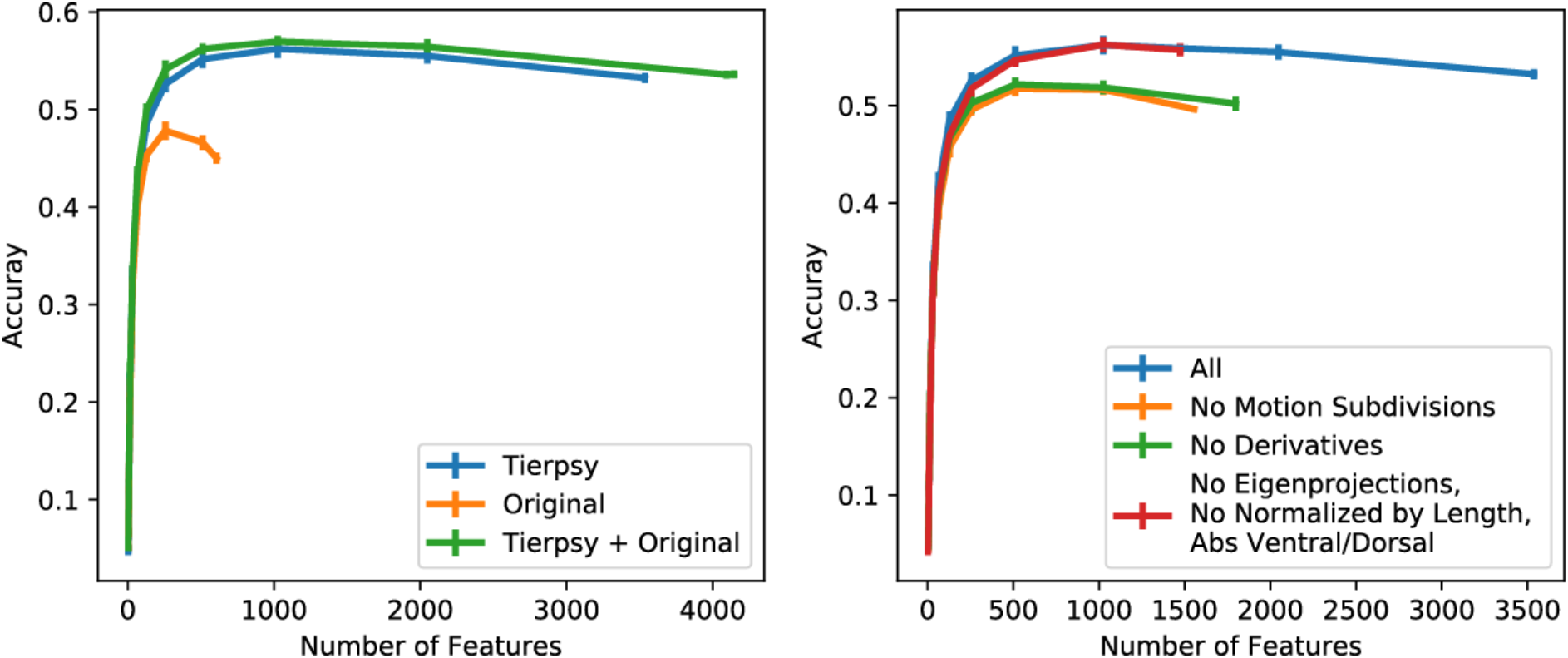
Results of recursive feature elimination. Left, the features reported in this paper (Tierpsy) achieve a higher accuracy than the ones used in Yemini *et al* [20] (Original). Right, the addition of transformations using subdivisions by motion type and derivatives over time are necessary to achieve the high accuracy. On the other hand, the eigenprojections, the normalization by length, and the ventral/dorsal sign have little or no effect.

The Tierpsy features perform better than the Yemini *et al*. features (peak accuracy of 56.21% at 1024 features compared to 47.82% at 256 features). The combined feature set shows the best performance, but the improvement over the Tierpsy features is small. This suggests that the Tierpsy features capture almost all of the phenotypic information in the Yemini *et al*. features as well as some new information that was missed. There is no drop in performance (and even a slight increase) as the first several thousand features are eliminated. The shape of the accuracy curve is highly reproducible when different subsets of worms are used for classification while the identity of the most useful features is highly variable, suggesting that many of the features in the total set are redundant. That is, within a set of correlated features, it makes little difference to classification accuracy which feature is kept and which are dropped.

The features that are selected also depend on nature of the strains used for feature selection. For example, if only wild isolate strains (rather than the full set of mutants which contain several strains with severe locomotion defects) are used for feature selection, the performance on classification on the mutant data is reduced (Fig. S2). In contrast, if all strains or only mutants are used in feature selection, the performance on classifying wild isolates is unaffected. This may be because the nature of the variation between wild isolates is represented by the differences between some mutants whereas there are differences between mutants that are no observed in wild isolates. This supports the choice of using a set of strains with a broad range of phenotypes in feature selection if the goal is to find a feature set that has the best chance of generalising to unseen worm strains.

While classification accuracy does not allow us to prioritise features within correlated groups, interpretability can provide a guide. To bias the results towards simple interpretable features, we started by eliminating classes of features and steps in the feature expansion. We found that removing derivatives and the subdivision by motion state both significantly reduced classification accuracy confirming that these are useful operations (Fig. 3b). However, we found it was possible to eliminate eigenworm features and the normalization by worm length, and to use only the absolute value of features that had previously been signed as positive or negative based on dorso-ventral orientation (e.g. curvature was originally defined as positive or negative depending on whether the body bend was dorsal or ventral). Together, removing these features reduces the total number by almost half with no detectable effect on accuracy. We label this reduced set of features the Tierpsy_2k.

For accurate classification or for clustering applications where the full spectrum of differences and similarities would be useful, we recommend a reduced set of 256 features, which we label the Tierpsy_256 (Table S1), that balances completeness and compactness of the representation. Many phenotyping tasks occur on a smaller scale, with just a handful of strains compared to a reference (such as several mutants compared to a wild type strain). If there are specific hypotheses for relevant phenotypic differences, having a large number of features to choose from makes it more likely that the hypotheses will be testable without having to code new features. However, for exploratory work, 256 features can lead to a larger number of differences than are needed to guide experiments and the large number of features increases the burden of multiple testing corrections. We have therefore used a combination of classification power, subjective interpretability, and coverage of feature classes to define the Tierpsy_8 and Tierpsy_16 (Table 1), which give classification accuracies of 20.37% ± 0.41% and 28.67% ± 0.45% respectively (mean ± standard deviation).

**Table 1:** List of manually selected features. The first eight features correspond to Tierpsy_8, while the whole list corresponds to Tierpsy_16.

|  |
| --- |
| length_90th |
| width_midbody_norm_10th |
| curvature_hips_abs_90th |
| curvature_head_abs_90th |
| motion_mode_paused_fraction |
| motion_mode_paused_frequency |
| d_curvature_hips_abs_90th |
| d_curvature_head_abs_90th |
| width_head_base_norm_10th |
| motion_mode_backward_frequency |
| quirkiness_50th |
| minor_axis_50th |
| curvature_midbody_norm_abs_50th |
| relative_to_hips_radial_velocity_tail_tip_50th |
| relative_to_head_base_radial_velocity_head_tip_50th |
| relative_to_head_base_angular_velocity_head_tip_abs_90th |

Pre-selecting these smaller sets of features before performing a new analysis reduces the multiple testing burden and results in phenotypic fingerprints that can be visualised and understood at a glance (Fig. 4).

**Figure 4:**
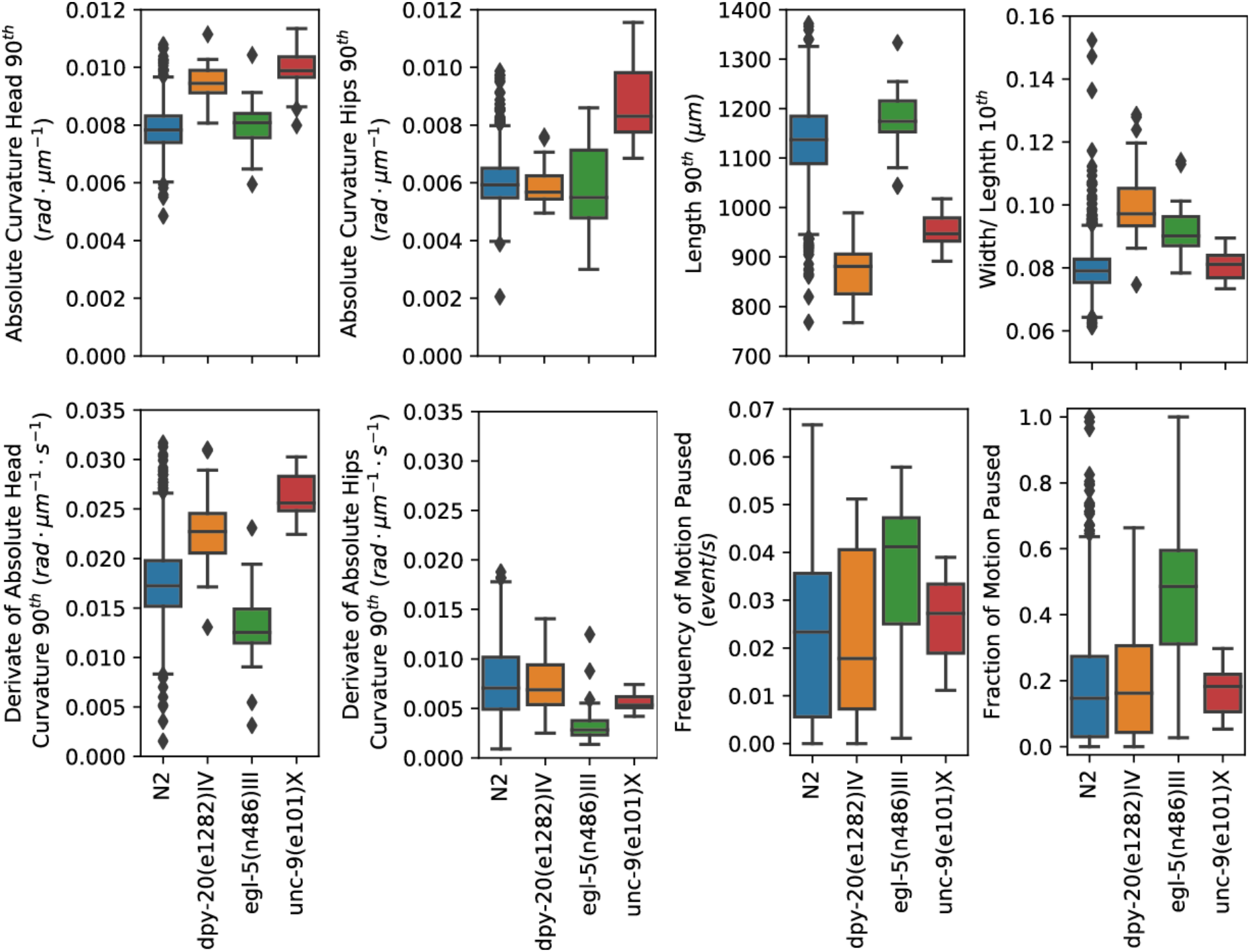
Boxplots of N2 and three mutants using the Tierpsy_8 subset of features. The small set of features facilitates a visual comparison between different strains.

### Direct analysis of time series for optogenetic experiments

The feature expansion and summarisation above captures some dynamic aspects of phenotype, but is intended for comparisons where the relevant differences are not localised in time and could occur at any point during a recording. For optogenetic experiments where the stimulation has a clear start and stop, it makes more sense to align time series and look for differences directly rather than summarising the entire experiment in a feature vector.

As above, we wanted to analyse data with a range of phenotypic differences. We therefore collected data from 11 strains expressing channel rhodopsin in different neural subsets (Table 2). We separate the data for each video into short pulses (five 5 second long segments) and long pulses (one 90 second long segment) and calculate histograms for each feature in each set. The behaviour may vary during or after a pulse and therefore it is useful to use multiple time bins to capture transient effects. We pooled the histograms for each strain with ATR and the controls without ATR. Because worms do not make ATR, any behavioural effects observed in the no-ATR condition are more likely to be generic blue light responses rather than ChR2-specific effects. The blue light response is most clearly observable during the 90 second pulses (Fig. S3).

**Table 2:** List of strains used in optogenetic experiments, their genotypes, and the neurons where channelrhodopsin is expressed.

| Strain | Neuron | Genotype (all found information) |
| --- | --- | --- |
| AQ2028 | CEP, ADE, PDE | lite-1(ce314); lJIs100 [dat-1::Chr2::YFP; Punc-122::GFP] |
| AQ2051 | M3R, M3L | lite-1 (ce314); Ex206 [eat-4::Chr2::YFP] |
| AQ2052 | ASH, ASI, PVQ | lite-1 (ce314); lJIs105 [sra-6::Chr2::YFP, Punc-122::GFP] |
| AQ2232 | AVM, ALM, PVM, PLM | lite-1 (ce314); lJIs111 [mec-4::Chr2] |
| AQ2235 | ADF, ASH, AWC, PHA, PHB | lite-1 (ce314); lJIs114 [sra-6::FTF::Chr2::YFP, gpa-13::FLPase] |
| AQ3071 | HSN, GLR | RQ10; wzIs6 [egl-6a::Chr2::YFP] |
| HBR180 | AVA, AVD, AVE, AVB, PVC | goeIs25 [nmr-1::Chr2::mKate2-unc-54-3'utr, unc 119 (+)] [suspected] |
| HBR187 | AVA, AVD, AVE, AVB, PVC | goeIs28 [nmr-1::Chr2::mKate2-unc-54-3'utr, unc 119 (+)] [suspected] |
| HBR222 | AVM, ALM, PVM, PLM | goeIs43 [mec-4::Chr2::mKate2-unc-54-3'utr, unc 119 (+)] |
| HBR520 | RIS | goeIs101 [aptf-1-5'utr::Chr2::mKate2-aptf-1-3'utr, unc119(+)] |
| MW544 | AWB | raxIs15 [str-1::Chr2::s12GFP] |

A selection of responses to 5 second optogenetic activation are shown in Fig. 5. There are clear differences between treatment and controls in several features for the three strains expressing ChR2 in different neural circuits. To systematically find features that respond differently to blue light between treatment and no-ATR controls, we calculated the Jensen-Shannon divergence between the treatment and control distributions for each feature and used a permutation test to determine a p-value for the comparison. Finally, we corrected for multiple comparisons within a given strain using the Benjamini-Hochberg procedure to control the false discovery rate at 0.05 [37]. The results are summarised in Fig. 6. Only five strains show p-values smaller than 0.05 after correction (AQ2235, AQ2052, AQ2232, HBR180, HBR520) in the short pulses (Fig 6 left).

**Figure 5:**
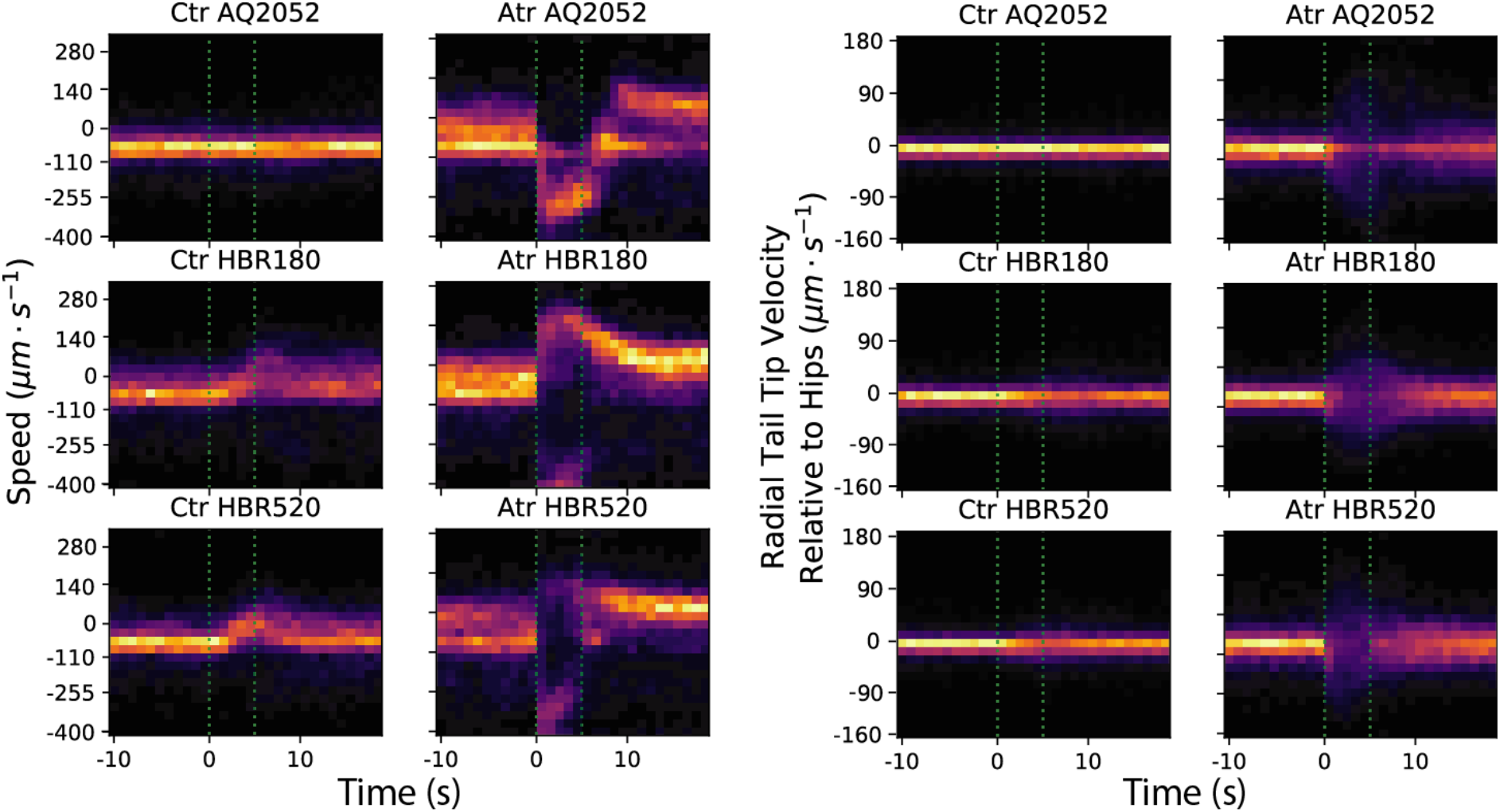
Example of 2D histograms of short pulses for different strains and features.

**Figure 6:**
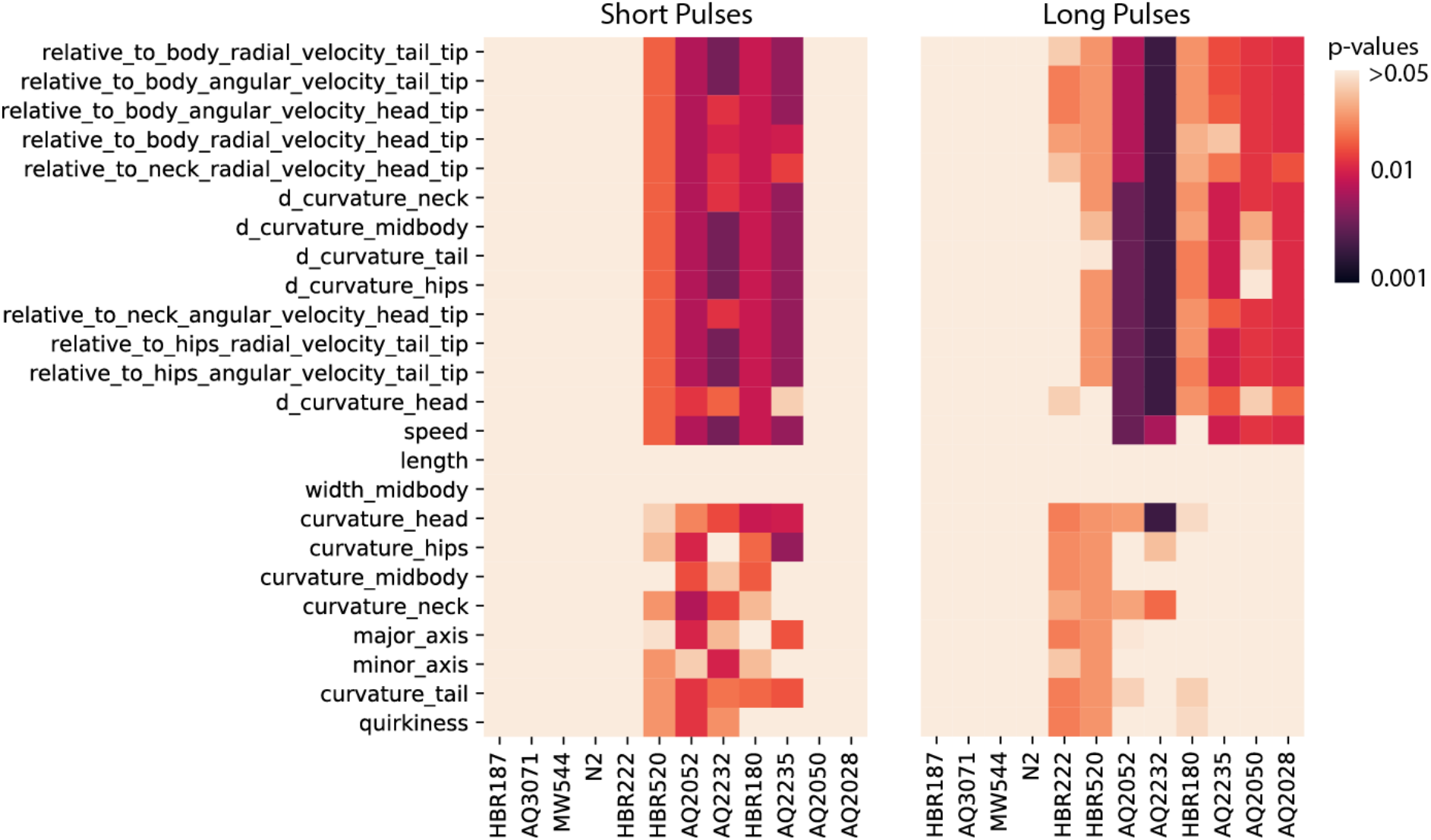
Comparisions between the changes of behaviour among the different strains under blue light stimulation for selected features. The heatmaps show the p-values of the Jensen-Shannon divergence between ATR and control plates for the short pulse experiments (left), and the long pulse experiments (right). The p-values were corrected for multiple comparisons within strains using the Benjamini-Hochberg procedure.

On the other hand, three additional strains show differences (HBR222, HBR520, AQ2050, AQ2028) when analysing the long pulse data (Fig 6 right), suggesting a slower response to optogenetic stimulation in these strains. Finally, it is worth noting that the three strains that did not show a significant difference compared to the control do not have a *lite-1* deletion mutation, so it is possible that an underlying optogenetic effect is masked by their aversive blue light response.

The permutation tests show that behaviour is significantly affected by optogenetic stimulation in several strains, but do not show how the features change to allow a comparison between strains. In order to quantitatively compare the behavioural responses between strains, we calculated a distance matrix between each strain in each condition (control and ATR) using the Jenson-Shannon divergence among the corresponding feature histograms. The results for the samples with ATR are shown in Fig 7. Two thirds of the strains are close to N2 in the long pulses but fewer than half are close to N2 in the short pulses. The observed clustering pattern suggests that there is a range of distinct behavioural phenotypes induced by the optogenetic stimulation of different neural subsets. The equivalent plots for the control plates are shown in Fig S4.

**Figure 7:**
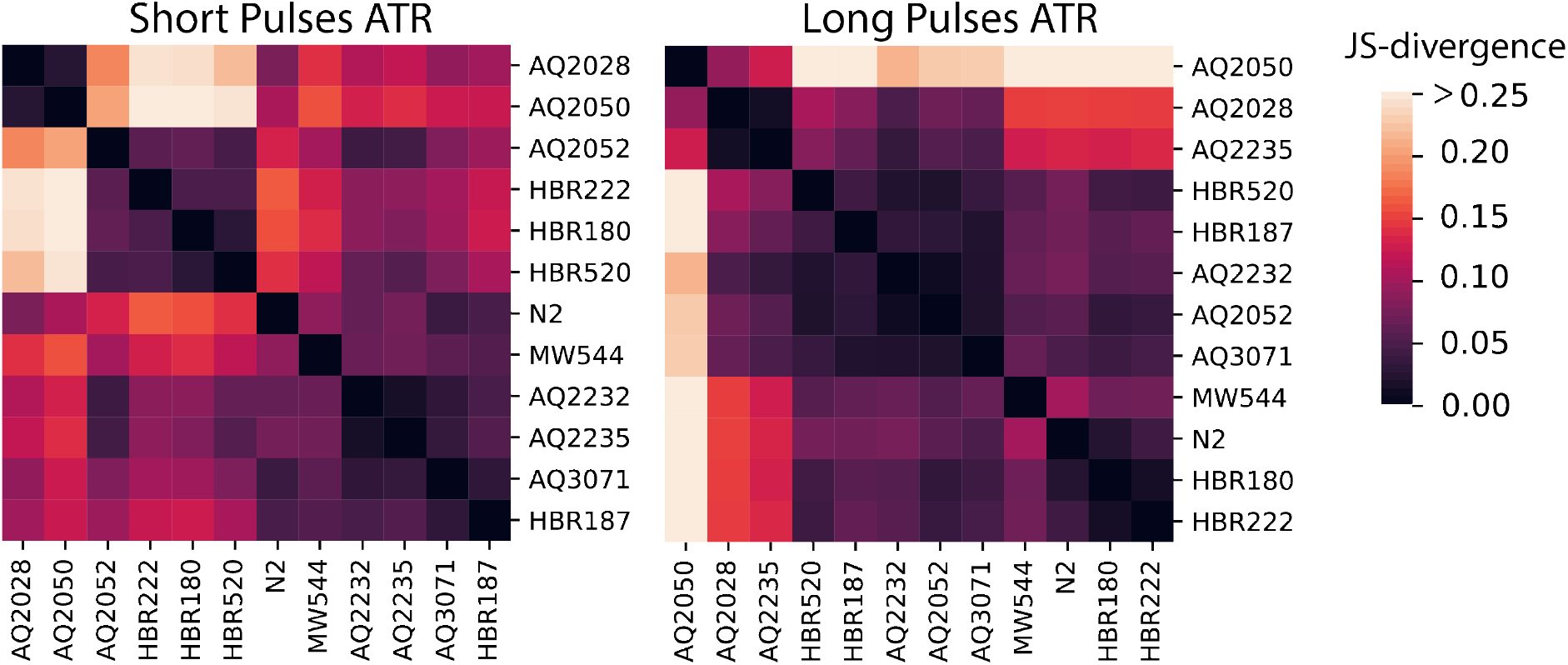
Cluster maps of the median value of the Jensen-Shannon divergence between different strains among the ATR plates.

## Discussion

The features we have defined here are intended to provide a range of options to balance power and interpretability in behaviour representation from a small set of easily interpretable features to a large set of features that provide improved classification accuracy compared a previously-defined set of handcrafted features. Using a diverse set of worm behaviour data from mutants and wild isolates, we found, as expected, that our feature definitions led to groups of correlated features that contained redundant information. We took advantage of this redundancy to favour interpretable features over equally useful but less interpretable features.

We found that two categories of features could be eliminated entirely with little effect on classification accuracy: those derived from the distribution of eigenworm amplitudes and those based on dorsoventral asymmetries. There is no contradiction between the elimination of eigenworm features and their usefulness in other applications [19,29,38,39]. The implication of this result is simply that the information present in the distribution of the eigenworm amplitudes taken across an entire video is captured using other more interpretable features. The eigenworm representation remains useful for many other applications, especially where an understanding of postural time series is important, rather than phenotypic summary based on posture distributions calculated for an entire video. Similarly, the known asymmetry between dorsal and ventral turns will clearly be important in some studies. It is just that on average, for distinguishing worm strains, it is not a critical distinction and most of the information is present in the symmetrised data. This is a positive result for multiworm tracking data where the dorsoventral orientation of worms is difficult to determine. Our results suggest that the Tierpsy features will be as useful for high resolution multi-worm tracking as they are for single worm tracking.

The data we used to perform feature selection covers a range of mutant phenotypes and the natural variation of wild isolates. Our goal was that the feature subsets we selected will be useful for capturing as complete a range of phenotypic variation as possible (Fig. S2). For applications where interpretability is paramount, they provide a useful starting point. However, for any new application with a different kind of phenotypic variation such as new mutants or worms in different experimental conditions, a different subset of features could be more appropriate. Therefore, for applications where there is sufficient data to use a training set on feature selection and a hold-out set for testing, we would recommend repeating the feature selection procedure starting from the full set of features or the Tierpsy_2k. Alternatively, if prior knowledge or specific hypotheses suggest a certain class of features is important, manually selected features can be simply added to one of the smaller feature subsets to capture the relevant effect without unduly increasing the burden of multiple testing.

## Conclusions

A critical step in any phenotyping project is choosing the right representation for the problem at hand. Phenomics is based on the assumption that the right representation can be difficult to determine *a priori* and that it is therefore useful to measure phenotypes as exhaustively as possible [1,2]. We have adopted this approach to define a large number of behavioural features that are then selected based on how well they explain data from a diverse set of strains and based on a subjective assessment of their interpretability. The direct parameterisation of behaviour we describe here is just one approach to the larger problem of the quantitative analysis of behaviour characteristic of computational ethology [40–42]. We have found that this approach has reasonably good power to detect subtle behavioural differences and is particularly useful in cases where interpretability is paramount.

In this paper we have focused on applications of phenotyping to analyse mutant strains, wild isolates, and optogenetically stimulated animals, but the same features could be useful in phenotypic drug screens, to quantify behavioural declines with ageing, or to characterise disease models.

One of the reasons the phenomic approach to behaviour analysis is useful is that animal behaviour is complex and the effects of perturbations are difficult to predict. The same is increasingly true for artificial agents controlled by artificial neural networks, which has led to calls for an ethology of artificial agents to understand their behaviour [43]. When applied to understanding the output of increasingly detailed simulations of *C. elegans* [44], we believe that a high-dimensional representation of behaviour will be essential to provide enough constraints to fit model parameters that are difficult to estimate directly from experiments.

We propose that the features defined here are a useful starting point for performing quantitative model validation for cellular level simulations based on the *C. elegans* connectome.

## Acknowledgements

This work was supported by grants to AEXB from the European Research Council (PHENOSPACE 714853) and Medical Research Council (MC-A658-5TY30). Some strains were provided by the CGC, which is funded by NIH Office of Research Infrastructure Programs (P40 OD010440). We thank Bill Schafer for the gift of strains AQ2028, AQ2051, AQ2052, AQ2232, AQ2235, and AQ3071, Henrik Bringmann for the gift of strains HBR180, HBR187, HBR222, and HBR520, and Meng Wang for strain MW544.

## Features Definitions

### Body Parts

The default skeleton is contains 49 points equally distributed along the length of the worm body. If the number of segments is different, the corresponding indices will be calculated as 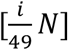. The segment limits of each body part are shown in the table below.

| Part | Lower Index | Upper Index |
| --- | --- | --- |
| Head | 1 | 8 |
| Body | 9 | 41 |
| Tail | 42 | 49 |
| Head Tip | 1 | 3 |
| Head Base | 4 | 8 |
| Neck | 9 | 16 |
| Midbody | 17 | 33 |
| Hips | 34 | 41 |
| Tail Base | 42 | 44 |
| Tail Tip | 45 | 49 |

### Morphology

#### Length

The length of a skeleton with *n* points *p* is defined as the sum of all the euclidean distances between two consecutive points in the skeleton as:

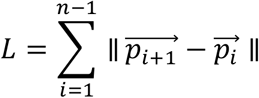

#### Area

The area is calculated from the worm contour using the shoelace method defined as:

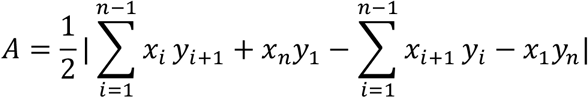

where *x* and *y* are the coordinates of a non-crossing contour with *n* elements. This method will give incorrect results for coiled (self-intersecting) worms shapes, but these are not currently skeletonised by the current algorithm.

#### Widths

Median contour width at specific parts of the body. The width at each point of the skeleton must be given as an input by the user (this is calculated in an earlier processing step by Tierpsy Tracker).

### Postures

#### Curvature

The curvature *κ* is defined as the rate of change of a curve’s tangent angle with respect to its arclength and is given by

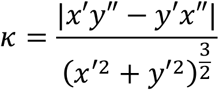

where *x* and *y* are the coordinates of each point *p*. We calculate the numerical derivatives as

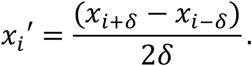

In order to obtain a smoother gradient, we use a *δ* = 2.

Finally, for each curvature along the skeleton we summarize the data using:

- The curvature at specific points in the curve.
- The mean and the standard deviation along specific body part ranges.

#### Major axis, minor axis and quirkiness

We use the OpenCV function *minAreaRect* to calculate the minimum-area bounding rectangle, and define the smallest and largest sides of the rectangle as the minor *a* and major *A* axis respectively.

We define quirkiness as

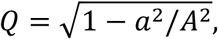

making it a metric analogous of the eccentricity but using the bounding box axes instead of the ones of a fitted ellipse.

#### Eigenworms coefficients

Following the procedure of Stephens *et al*. [1], the frame of reference is changed by calculating the tangent angle between consecutive points as

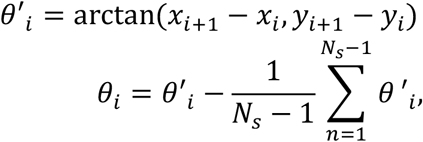

where *N_s_* is the total number of segments in the skeletons, *x_i_* and *y_i_* are the segment coordinates, and *θ_i_* is the corresponding segment angle. The final features are coefficients of the angles projected onto the eigenworms as

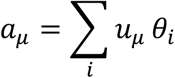

where *μ* is the eigenworm number and *u* are the eigenworms previously calculated using principal components analysis on the data in Yemini *et al*. [2]. We use the first 7 eigenworms, which capture more than 98% of the variance in the original dataset.

### Velocities

All the velocities are calculated using a user defined time window *Δt* with a default value of 1/3*s*.

#### Speed and Angular Velocity

For each body part segment, we calculate its centroid

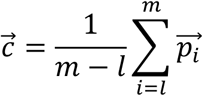

and its orientation vector

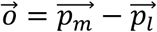

where *l* and *m* are respectively the lower and upper indices of a given body part, and 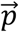 are the coordinates of the skeleton.

The angular velocity is calculated as

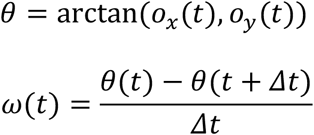

For the speed, we first calculate the centroid *c* velocity as

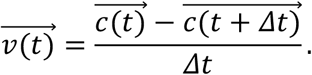

and then calculate the signed speed as

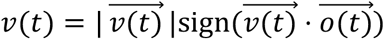

so the speed is positive when the worm is moving forwards and negative when it is moving backwards.

#### Relative Velocities

The centres of mass are calculated as above. We then shift the coordinate system of a given body part 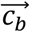 to be centred with respect to a reference body part 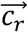 as

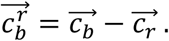

The angular and radial velocity are calculated by changing from the cartesian (*x*, *y*) to the polar coordinate system (*ϕ, r*). The relative angular velocity is the derivative with respect to time of the *ϕ* coordinate of 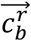, while the relative radial velocity is the same but for coordinate *r*.

The relative midbody velocity is a special case where *b* is the midbody and *r* is the body, and the time derivative is calculated with respect to the cartesian coordinate *x*.

### Path

The path features with respect to a given body part are calculated over the centre of mass of the corresponding skeleton segment as above.

#### Path Curvature

We resample the path over space so each point in the trajectory is equidistant, and then calculate the curvature equation described in the Curvature section. Finally, we interpolate curvature back so each element is equally separated in time.

#### Path Grid

The path grid is a 2D histogram where each cell represents a small region in the recording area. The counts represent the amount of time (in frames) spent in a given cell. Each side of the individual cells has a default value 250*μm*. From this grid the following features are extracted:

***Path Coverage*** is total area explored by the worm, calculated as the total number of grids explored by the worm times the individual grid area.
***Path Density*** is the probability of a worm staying in a given cell calculated as the counts per cell over the total number of counts. Only the 50th and 90th percentiles of the density distributions are calculated.
***Grid Transit Time*** is the time spent in a cell before moving to the next one. Only the 50th and 90th percentiles of the transit time distributions are calculated.

### Transformations

***Time derivative*** of a given time series feature are calculated over a window with a default value of 1/3s, similar to the velocity.

***Percentiles:*** 10th, 50th and 90th percentiles of a given time series feature.

***Interquantile Range (IQR):*** calculated as the 75th minus the 25th percentile of a given time series distribution. It is a robust statistic of the distribution spread.

***Normalization by Worm Length:*** the units of a given feature are converted from *μm* to body lengths.

***Absolute Ventral/Dorsal Sign:*** the value of a feature is signed according to the ventral/dorsal orientation [2].

***Subdivision According to Movement:*** a time series is subdivided into segments where the worm is moving either forward, backwards, or is paused. These movement states are determined from the worm velocity.

---

**Figure S1:**
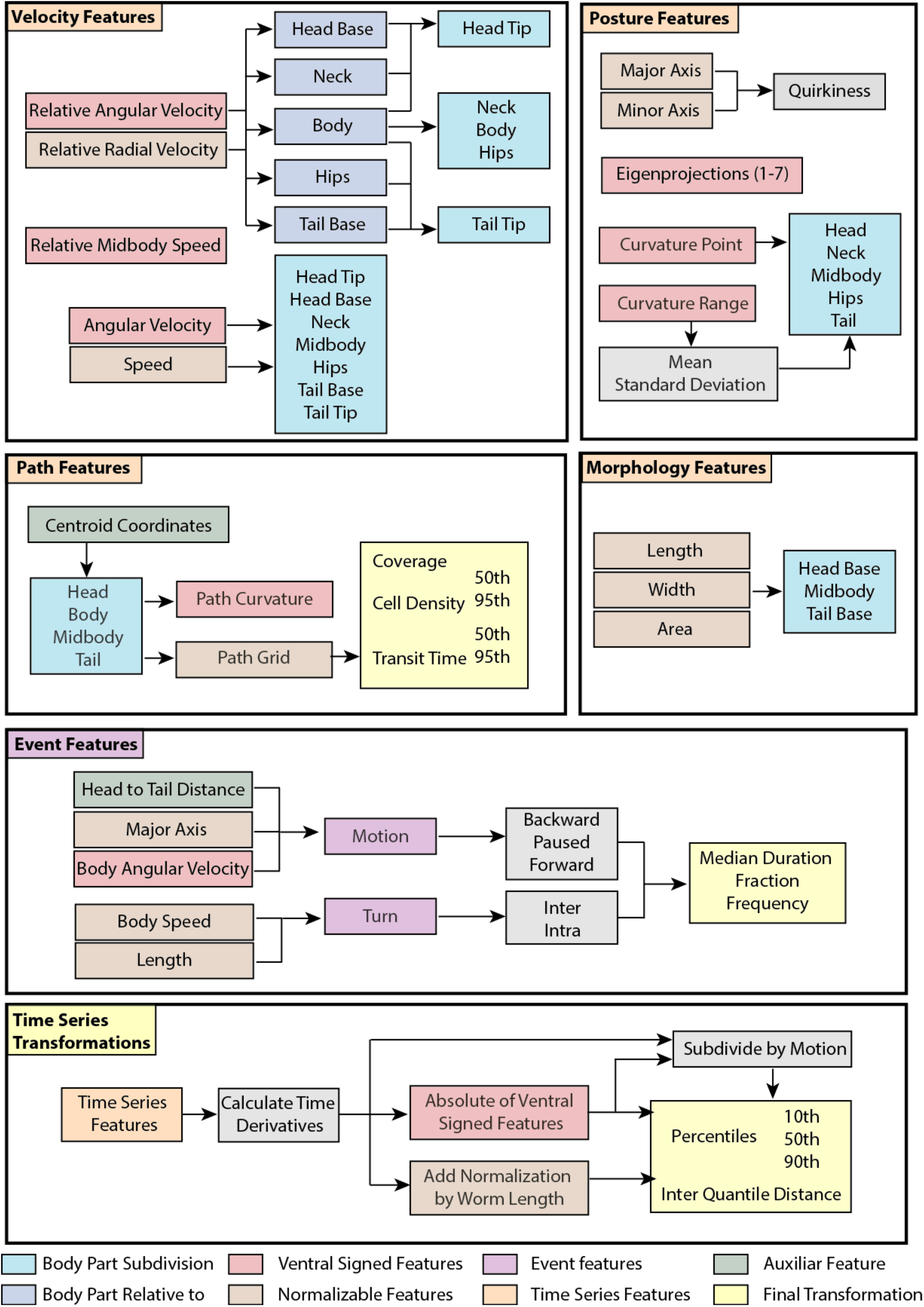
Operations that expand and summarize each of the core features.

**Fig. S2:**
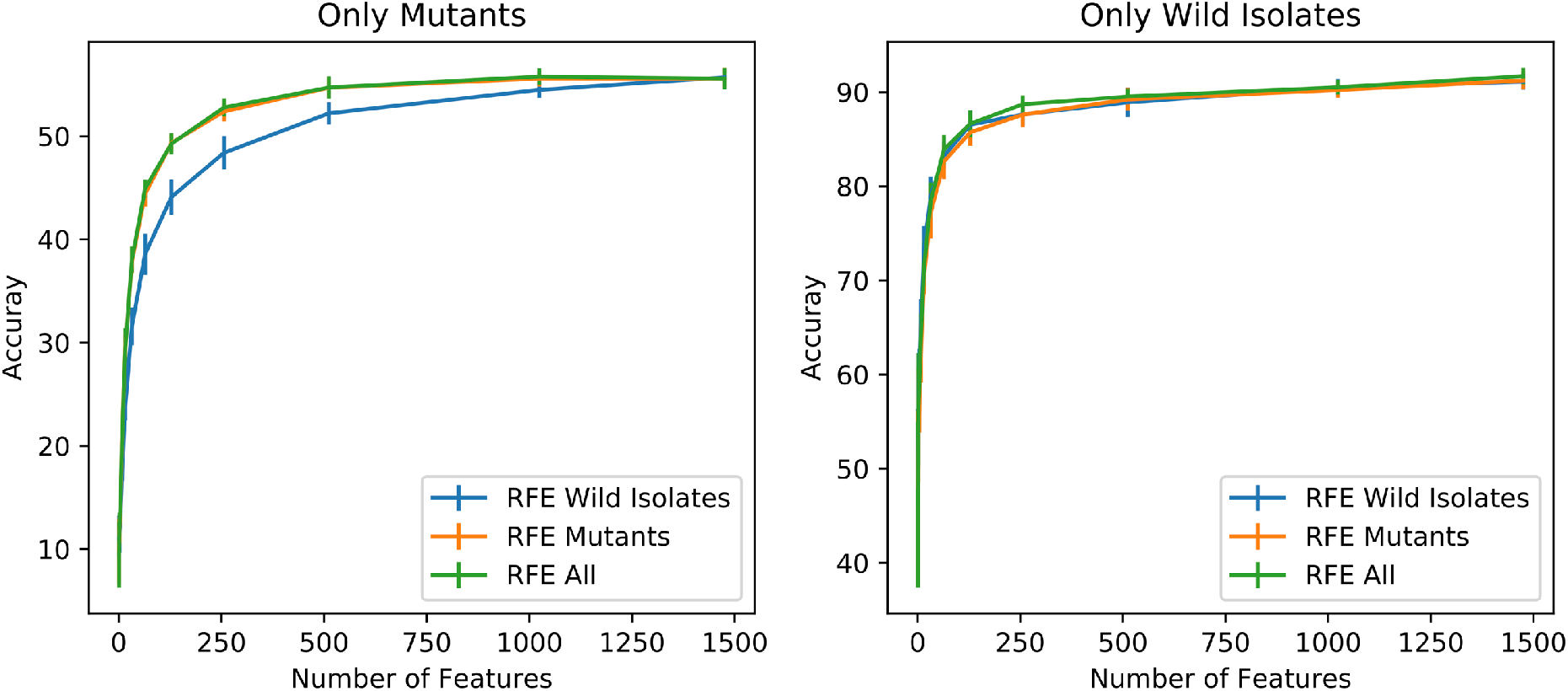
Comparison of classification performance using different strain subsets for feature selection using recursive feature elimination (RFE). Each curve shows the results using features selected using all strains, mutant strains without wild isolates, or wild isolates only. The plot on the left shows the classification performance of each feature subset on the mutant data. The plot on the right shows the classification performance of each feature subset on the wild isolate data. The features selected using only wild isolate data perform worse on classifying mutants, while the features selected using mutants perform as well as features selected directly on wild isolates for classifying wild isolates.

**Figure S3:**
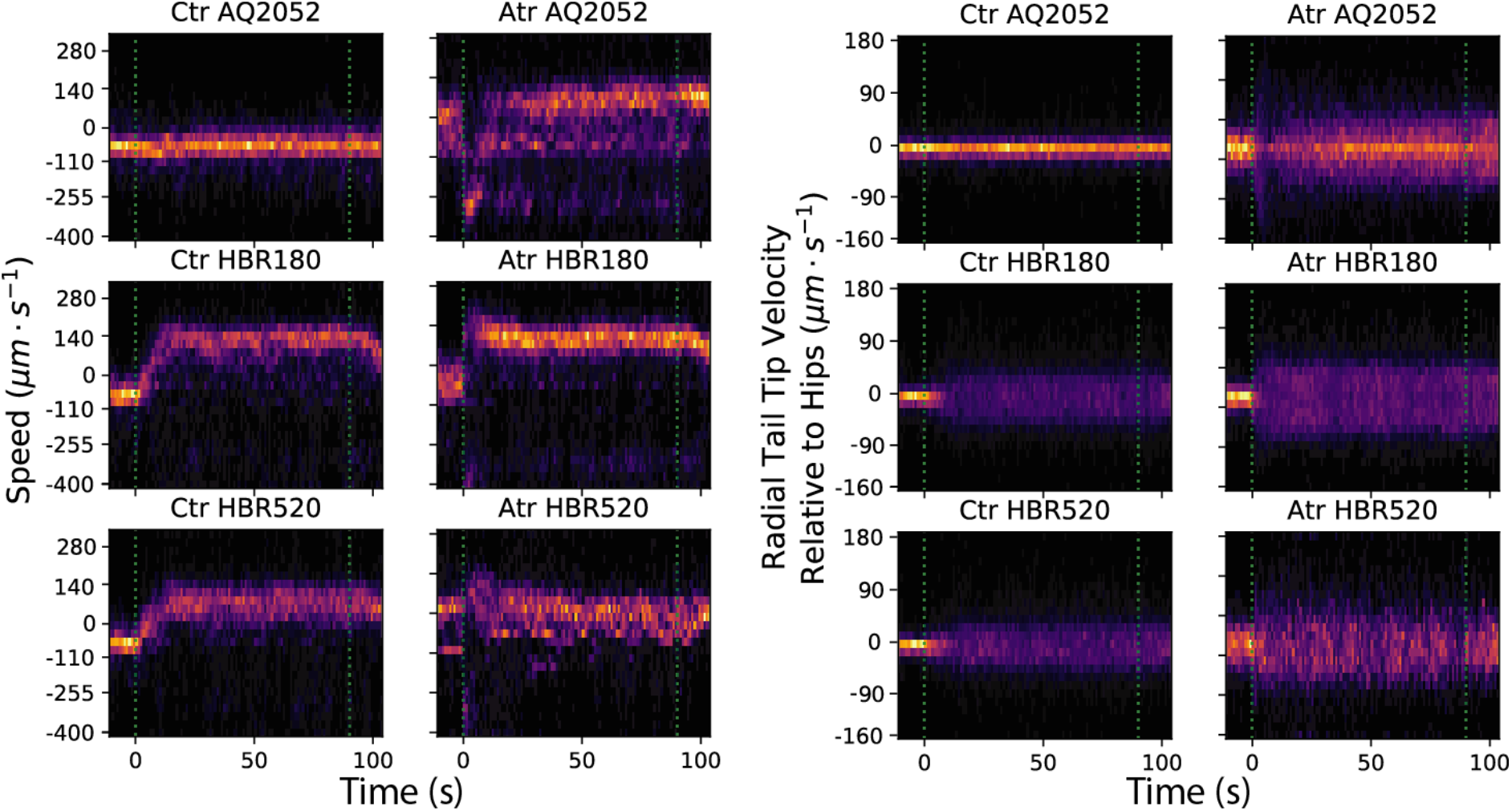
Example of 2D histograms of long pulses for different strains and features.

**Figure S4:**
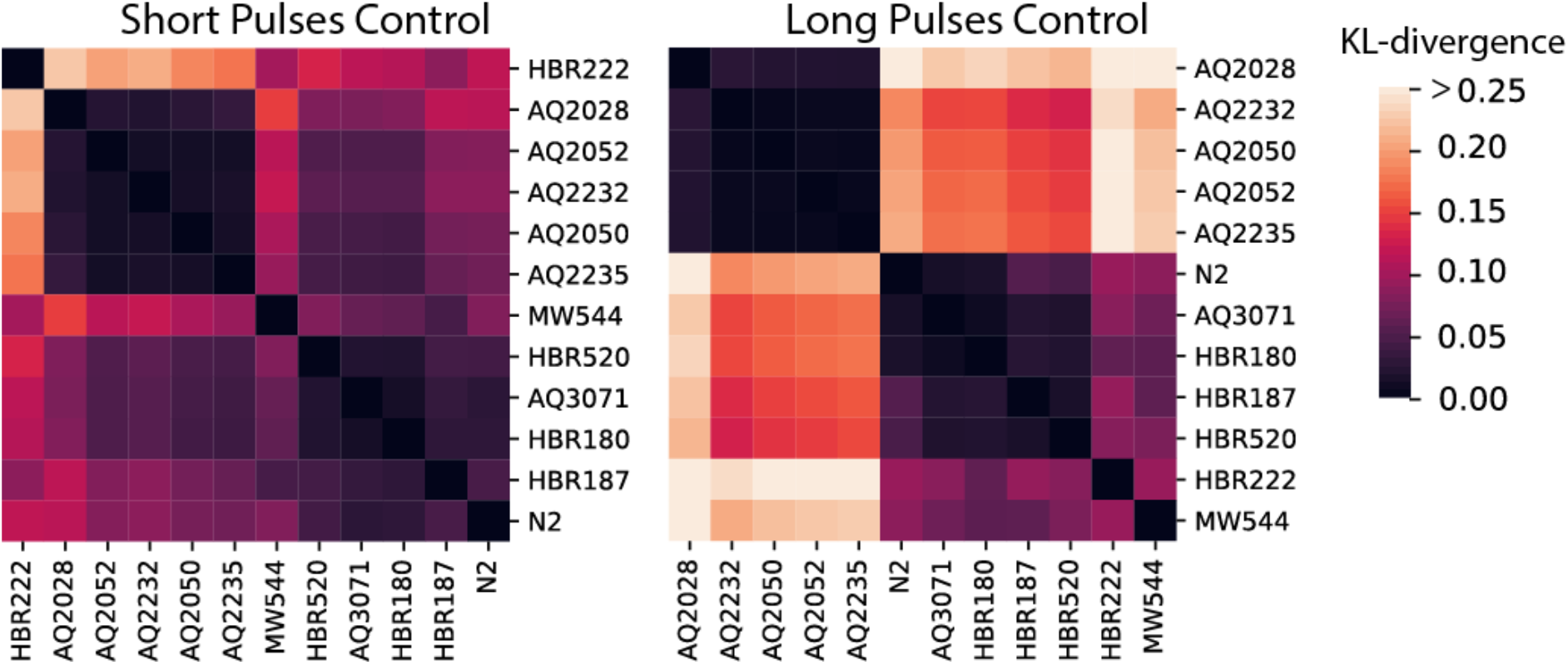
Cluster maps of the median value of Jensen-Shannon divergence between different strains for the control plates.

**Table S1:** There are many sets of 256 features that perform similarly in classifying the strains used in this paper. For this larger set, we have not applied an interpretability criterion. Nonetheless, for concreteness and reproducibility, we include one instance, which we label the Tierpsy_256 here.

|  |
| --- |
| motion_mode_paused_frequency |
| motion_mode_paused_fraction |
| curvature_std_neck_abs_90th |
| d_curvature_neck_w_backward_abs_90th |
| angular_velocity_tail_tip_w_backward_abs_IQR |
| curvature_std_midbody_abs_10th |
| relative_to_head_base_radial_velocity_head_tip_50th |
| width_tail_base_w_forward_50th |
| minor_axis_w_forward_10th |
| d_relative_to_head_base_angular_velocity_head_tip_w_backward_abs_IQR |
| d_relative_to_body_radial_velocity_head_tip_w_forward_90th |
| relative_to_head_base_angular_velocity_head_tip_abs_50th |
| relative_to_body_radial_velocity_tail_tip_w_backward_10th |
| d_angular_velocity_tail_base_w_backward_abs_90th |
| relative_to_body_radial_velocity_hips_w_forward_90th |
| width_head_base_w_forward_10th |
| curvature_std_tail_w_forward_abs_90th |
| d_area_w_backward_10th |
| d_curvature_mean_hips_w_forward_abs_IQR |
| relative_to_head_base_radial_velocity_head_tip_w_forward_10th |
| width_midbody_10th |
| curvature_head_abs_90th |
| d_curvature_std_head_w_forward_abs_50th |
| curvature_mean_head_abs_90th |
| curvature_std_neck_w_forward_abs_50th |
| d_width_head_base_IQR |
| d_minor_axis_w_backward_90th |
| length_w_backward_10th |
| width_midbody_w_forward_10th |
| motion_mode_forward_fraction |
| curvature_std_head_w_forward_abs_50th |
| relative_to_neck_radial_velocity_head_tip_50th |
| motion_mode_backward_frequency |
| curvature_std_neck_abs_10th |
| angular_velocity_abs_IQR |
| length_IQR |
| d_angular_velocity_head_tip_abs_10th |
| relative_to_hips_radial_velocity_tail_tip_50th |
| d_speed_head_tip_50th |
| relative_to_head_base_radial_velocity_head_tip_w_forward_90th |
| quirkiness_50th |
| relative_to_head_base_radial_velocity_head_tip_w_backward_90th |
| d_width_tail_base_IQR |
| minor_axis_w_forward_90th |
| d_relative_to_neck_radial_velocity_head_tip_w_forward_90th |
| d_relative_to_tail_base_radial_velocity_tail_tip_50th |
| curvature_head_w_forward_abs_IQR |
| relative_to_head_base_radial_velocity_head_tip_w_backward_IQR |
| d_path_curvature_midbody_abs_90th |
| relative_to_body_radial_velocity_head_tip_w_forward_50th |
| d relative to head base radial velocity head tip w backward 50th |
| d length 50th |
| width_midbody 90th |
| curvature_hips w backward abs 50th |
| path_transit_time_body_95th |
| d_length_w_forward_IQR |
| motion_mode_backward_fraction |
| speed_head_tip_w_forward_IQR |
| curvature_std_hips_abs_10th |
| relative_to_head_base_angular_velocity_head_tip_abs_IQR |
| d_major_axis_w_backward_50th |
| d_curvature_std_tail_w_forward_abs_90th |
| curvature_mean_hips_w_backward_abs_90th |
| path_coverage_head |
| d_relative_to_tail_base_angular_velocity_tail_tip_w_forward_abs_90th |
| d_relative_to_head_base_angular_velocity_head_tip_w_backward_abs_90th |
| relative_to_head_base_radial_velocity_head_tip_w_forward_IQR |
| width_tail_base_w_backward_IQR |
| curvature_mean_neck_abs_50th |
| curvature_std_neck_w_forward_abs_10th |
| curvature_std_midbody_w_backward_abs_10th |
| d_major_axis_50th |
| curvature_hips_abs_90th |
| relative_to_body_radial_velocity_head_tip_50th |
| curvature_tail_w_backward_abs_90th |
| curvature_mean_head_w_forward_abs_50th |
| d_width_midbody_w_forward_10th |
| width_midbody_w_backward_50th |
| relative_to_head_base_radial_velocity_head_tip_10th |
| curvature_midbody_abs_90th |
| curvature_tail_abs_90th |
| curvature_mean_neck_w_forward_abs_IQR |
| width_tail_base_w_backward_10th |
| turn_intra_duration_50th |
| d_relative_to_head_base_angular_velocity_head_tip_w_forward_abs_90th |
| width_midbody_w_backward_10th |
| d_curvature_std_head_w_forward_abs_10th |
| curvature_mean_tail_abs_90th |
| speed_10th |
| d_angular_velocity_head_tip_w_forward_abs_90th |
| relative_to_body_radial_velocity_hips_50th |
| d_width_tail_base_50th |
| width_tail_base_10th |
| d_speed_head_tip_w_forward_50th |
| d_angular_velocity_head_tip_w_forward_abs_50th |
| d_width_head_base_w_backward_50th |
| speed_head_tip_10th |
| curvature_std_head_abs_IQR |
| curvature_hips_w_backward_abs_IQR |
| d_width_midbody_50th |
| d_speed_head_tip_w_backward_50th |
| length_w_forward_IQR |
| width_tail_base_w_backward_50th |
| curvature_tail_abs_10th |
| d_width_head_base_w_forward_IQR |
| d curvature std midbody w backward abs 10th |
| d width head base w forward 50th |
| width head base 10th |
| d length w backward IQR |
| d curvature std hips w forward abs 90th |
| motion mode backward duration 50th |
| d curvature std head abs 90th |
| d length w backward 10th |
| curvature std midbody w backward abs 50th |
| curvature std midbody w backward abs 90th |
| relative to neck radial velocity head tip w forward 10th |
| curvature std hips w backward abs IQR |
| d path curvature tail w backward abs 90th |
| curvature head w forward abs 50th |
| relative to body radial velocity neck 50th |
| relative to neck radial velocity head tip w forward 50th |
| d relative to body radial velocity tail tip w backward 10th |
| width head base w forward 50th |
| curvature mean tail w backward abs 90th |
| curvature midbody w forward abs 50th |
| d relative to head base radial velocity head tip 50th |
| length w backward IQR |
| d relative to neck radial velocity head tip w forward 50th |
| d angular velocity midbody w backward abs 90th |
| d curvature std head w backward abs 50th |
| curvature std hips abs 50th |
| width tail base 90th |
| relative to body radial velocity tail tip 50th |
| angular velocity head base abs 10th |
| curvature std hips w backward abs 90th |
| d relative to head base radial velocity head tip w backward 10th |
| path curvature body abs 10th |
| speed w forward 90th |
| d relative to body radial velocity neck w forward 90th |
| d relative to body radial velocity tail tip w backward 50th |
| d angular velocity hips w backward abs 90th |
| d relative to body angular velocity hips w forward abs 10th |
| width midbody w forward 50th |
| d curvature mean tail w forward abs 10th |
| d quirkiness 50th |
| speed hips w backward 90th |
| d curvature hips w backward abs 90th |
| curvature mean tail w backward abs IQR |
| d curvature mean hips w backward abs 90th |
| angular velocity tail base w backward abs 90th |
| d width head base 50th |
| path transit time midbody 95th |
| path curvature midbody w backward abs 10th |
| speed w forward IQR |
| width midbody 50th |
| d curvature neck w backward abs 10th |
| d curvature mean neck w backward abs 90th |
| relative to hips radial velocity tail tip w forward IQR |
| angular velocity head tip w forward abs 50th |
| d curvature mean head w forward abs 10th |
| curvature_std_neck_abs_50th |
| angular_velocity_head_base_abs_50th |
| curvature_std_midbody_w_backward_abs_IQR |
| d_curvature_std_neck_w_backward_abs_10th |
| speed_tail_base_w_forward_IQR |
| angular_velocity_tail_base_w_forward_abs_50th |
| d_relative_to_body_radial_velocity_tail_tip_w_forward_IQR |
| relative_to_tail_base_radial_velocity_tail_tip_50th |
| d_angular_velocity_midbody_w_backward_abs_50th |
| curvature_std_neck_w_backward_abs_50th |
| width_midbody_w_backward_90th |
| d_relative_to_body_radial_velocity_hips_50th |
| curvature_std_neck_w_backward_abs_10th |
| d_relative_to_body_radial_velocity_tail_tip_w_forward_90th |
| d_curvature_head_w_backward_abs_50th |
| angular_velocity_head_base_w_forward_abs_10th |
| d_relative_to_body_angular_velocity_neck_w_backward_abs_10th |
| d_relative_to_body_angular_velocity_tail_tip_abs_10th |
| d_curvature_mean_midbody_abs_90th |
| width_tail_base_w_backward_90th |
| curvature_std_midbody_abs_90th |
| d_curvature_midbody_w_backward_abs_10th |
| curvature_tail_w_forward_abs_90th |
| curvature_std_midbody_abs_50th |
| angular_velocity_head_base_w_forward_abs_50th |
| speed_head_tip_w_forward_50th |
| d_curvature_head_w_forward_abs_10th |
| angular_velocity_head_tip_w_backward_abs_IQR |
| d_curvature_hips_w_backward_abs_10th |
| d_relative_to_body_radial_velocity_neck_w_forward_10th |
| d_relative_to_tail_base_radial_velocity_tail_tip_w_forward_10th |
| minor_axis_10th |
| speed_head_tip_w_forward_10th |
| d_relative_to_body_angular_velocity_hips_w_backward_abs_90th |
| d_curvature_mean_hips_abs_IQR |
| relative_to_tail_base_radial_velocity_tail_tip_90th |
| relative_to_neck_radial_velocity_head_tip_w_forward_90th |
| d_curvature_mean_hips_abs_90th |
| curvature_hips_abs_10th |
| d_major_axis_w_forward_IQR |
| d_angular_velocity_head_base_w_forward_abs_90th |
| curvature_std_midbody_abs_IQR |
| relative_to_tail_base_radial_velocity_tail_tip_w_backward_90th |
| curvature_midbody_abs_IQR |
| speed_head_tip_w_backward_10th |
| curvature_std_head_abs_90th |
| width_midbody_w_forward_90th |
| d_angular_velocity_midbody_w_backward_abs_IQR |
| d_curvature_hips_abs_90th |
| d_length_w_backward_90th |
| d_curvature_mean_head_w_backward_abs_50th |
| relative_to_body_radial_velocity_head_tip_w_backward_50th |
| relative_to_neck_radial_velocity_head_tip_w_forward_IQR |
| d_curvature_mean_head_w_forward_abs_50th |
| d_angular_velocity_abs_10th |
| angular_velocity_head_base_w_backward_abs_90th |
| d_major_axis_w_forward_10th |
| curvature_tail_w_forward_abs_10th |
| d_length_10th |
| d_curvature_std_hips_w_backward_abs_IQR |
| d_angular_velocity_head_tip_abs_90th |
| d_length_90th |
| d_angular_velocity_tail_base_w_backward_abs_50th |
| d_curvature_std_head_abs_IQR |
| curvature_mean_hips_w_backward_abs_IQR |
| speed_tail_base_w_backward_90th |
| angular_velocity_head_base_w_backward_abs_IQR |
| relative_to_neck_radial_velocity_head_tip_w_backward_50th |
| d_minor_axis_50th |
| d_relative_to_tail_base_radial_velocity_tail_tip_w_backward_90th |
| angular_velocity_head_tip_abs_50th |
| motion_mode_forward_frequency |
| width_tail_base_50th |
| d_curvature_mean_neck_w_forward_abs_IQR |
| angular_velocity_tail_base_w_forward_abs_IQR |
| curvature_std_head_abs_50th |
| speed_hips_w_forward_IQR |
| d_angular_velocity_tail_base_w_backward_abs_IQR |
| d_relative_to_body_radial_velocity_head_tip_w_backward_10th |
| d_relative_to_body_radial_velocity_tail_tip_w_forward_10th |
| d_curvature_mean_neck_w_backward_abs_10th |
| d_relative_to_head_base_radial_velocity_head_tip_w_forward_IQR |
| relative_to_neck_angular_velocity_head_tip_w_backward_abs_10th |
| relative_to_head_base_radial_velocity_head_tip_w_backward_10th |
| d_length_w_backward_50th |
| curvature_hips_abs_IQR |
| d_curvature_std_neck_w_backward_abs_50th |
| d_angular_velocity_neck_w_forward_abs_10th |
| d_speed_neck_50th |
| d_relative_to_body_radial_velocity_neck_w_backward_50th |
| d_major_axis_w_backward_10th |
| d_curvature_std_head_w_backward_abs_10th |
| d_path_curvature_midbody_w_backward_abs_90th |
| d_relative_to_body_angular_velocity_neck_w_backward_abs_90th |
| d_relative_to_hips_radial_velocity_tail_tip_50th |
| width_tail_base_w_forward_10th |

